# Functional Enrichment and Analysis of Antigen-Specific Memory B Cell Antibody Repertoires in PBMCs

**DOI:** 10.1101/557769

**Authors:** Eric Waltari, Aaron McGeever, Peter S. Kim, Krista M. M^c^Cutcheon

**Author notes:** Correspondence: Krista M. M^c^Cutcheon.

## Abstract

Phenotypic screening of antigen-specific antibodies in human blood is a common diagnostic test for infectious agents and a correlate of protection after vaccination. In addition to long-lived antibody secreting plasma cells residing in the bone marrow, memory B cells are a latent source of antigen-experienced, long-term immunity that can be found at low frequencies in circulating PBMCs. Assessing the genotype, clonal frequency, quality, and function of antibodies resulting from an individual’s persistent memory B cell repertoire can help inform the success or failure of immune protection. We have applied ELISPOT cell culture methods to functionally expand the memory repertoire from PBMCs and clonally map monoclonal antibodies from this population. We show that combining deep sequencing of stimulated memory B cell repertoires with retrieving single antigen-specific cells is a promising approach in evaluating the latent, functional B cell memory in PBMCs.

## INTRODUCTION

One of public health’s most cost-effective interventions to prevent and reduce the disease burden of infectious disease is vaccination. It is well established that immunological memory stored in antigen-specific B cell repertoires frequently forms the foundation of successful natural or vaccine-induced immune protection (Sallusto, F. *et al* 2010). During the acute phase of pathogen exposure, naïve B cell clones enter germinal centers and undergo Ig receptor somatic hypermutation (SHM) to increase antigen binding, isotype class switching to IgG or IgA for effector activities, and differentiation to short-lived antibody secreting plasmablasts or to long-lived plasma and memory B cells (De Silva, N.S. and Klein, U., 2015). Successful diversification of B cell clones and their corresponding Ig receptors creates clonal families of these cells with variations in antigen binding and functional activities. Antibodies secreted from plasma cells, which mainly reside in the bone marrow, provide steady state protection against repeat infections. In contrast, memory B cells survive in a functionally quiescent state in tissues and can be found at low frequency in PMBCs after a pathogen is eliminated, re-activating and differentiating into antibody secreting plasmablasts within a week of a secondary infection (Abbas A.K. *et al* 2018, Yoshida, T. *et al* 2010, Antia A. *et al* 2018). Evaluating both types of adaptive B-cell memory is challenged by the anatomical inaccessibility and rarity of these cells. A variety of serological assays for polyclonal antibody measurements have been developed (Madore, D.V. *et al* 2010) and, more recently, the applications of next generation sequencing (NGS) and single cell sequencing technologies have offered high resolution data on B cell antibody responses in peripheral blood samples (Laserson, U. *et al* 2014). Computational algorithms to interpret genetic relationships between the millions of NGS B cell receptor (BCR) sequences have been developed to assign clonal families (BCR deriving from a common B cell ancestor, sometimes referred to as clonotypes) and help reconstruct *in vivo* B cell clonal lineages (Gupta N.T. et al 2017). However, outside of the acute phase of exposure, the depth of antigen-experienced B cell repertoires is still limited by the low frequency and transcriptional silence of memory B cells circulating in human blood.

Herein, we show the benefit of combining NGS with *in vitro* cell culture methods, originally established for ELISPOT, that functionally and selectively expand memory B cells within PBMC samples using polyclonal stimulation with cytokines and a Toll-like receptor 9 agonist, CpG (Crotty S. *et al* 2004, Weiss G.E. *et al* 2012, McCutcheon K.M. *et al* 2014). Using three healthy donors of different genetic backgrounds we compared NGS data of immunoglobulins on PBMC repertoires with stimulated PBMC repertoires. In stimulated PBMCs we observed an enrichment of sequences of IgG subtypes and higher rates of SHM with no resulting bias in the representation of heavy or light chain variable domain families. We identified common V-D-J BCR sequences (clonal families) within one donor that persisted over three blood draws, spanning a 9-month period, in PBMC and in stimulated PBMC immunoglobulin repertoires. The majority of the persistent clones identified in the stimulated repertoire, while fewer in number, showed >1% SHM and isotype switching to IgG, consistent with being part of a memory population. From the same donor, we cloned antigen-specific monoclonal antibodies (mAbs) from single memory B cells isolated by FACS with the hemagglutinin trimer of influenza A H1N1. Members of these mAbs clonal lineages were identified in PBMCs versus stimulated PBMCs repertoires, quantified, and tracked over the three blood draws. An increased sensitivity for detection of individual mAb clones along with larger, branching clonal families were obtained using data from stimulated PBMCs. Overall, our data support the application of stimulating PBMC samples to gain a deeper understanding of antigen-specific circulating memory B cell repertoires.

## MATERIALS AND METHODS

### Isolation of PBMCs

Leukapheresis was performed on three normal donors using Institutional Review Board (IRB)-approved consent forms and protocols by StemCell Technologies (Vancouver, BC). Approximately two to three blood volumes were processed using the Spectra Optia® Apheresis System to produce a full-sized Leuko Pak. Acid citrate dextrose, solution A (ACDA) was the anticoagulant. Donors included: 536, African American age 33, male, 82kg, 180cm, smoker, collected 11/15/2017, 5/24/2018 and 08/27/2018; 147, Caucasian female age 21, 56kg, 153cm, non-smoker, collected 08/21/2018; and 682, Hispanic male age 30, 74kg, 181cm, smoker, collected 08/16/2018. PBMCs were further enriched from the buffy coats of 50ml tubes with 25ml sterile Histopaque 1.077 (MilliporeSigma, Burlington, MA) layered with 25ml leukocytes diluted 1:1 in PBS (no Ca^++^/Mg^++^), after centrifuging at 400xg, 20°C, for 50min (Eppendorf Centrifuge 5810, with swinging bucket rotor S-4-104, brake off). PBMCs from each 50ml tube were washed by diluting three times with 50ml ambient temperature RPMI 1640 and centrifuged 200xg, 20°C, 10min. They were cryopreserved in 90% heat-inactivated FBS, 10% DMSO at 1×10^7^ cells per vial.

### Preparation of Growth-Arrested Feeder Cells

Human fibroblast cell line MRC-5 (CCL-171) was obtained from ATCC® (Manassas, VA) and grown in B cell growth media to 80% confluence before being treated for 4h with 5μg/ml mitomycin C (Tocris, R&D Systems). Monolayers of growth arrested cells were washed 3 times with PBS, harvested with trypsin, neutralized with growth media, washed 1x in growth media and finally cryopreserved using 10% DMSO, 30% HI-FBS in growth media.

### Preparation of PBMCs for BCR Repertoire Analysis

PBMC vials were thawed rapidly in a 37°C water bath, immediately diluted into 10ml of B cell growth media containing Corning® DMEM [+] 4.5 g/L glucose, sodium pyruvate [−] L-glutamine (VWR International, Radnor, PA), 1xPen/Strep/Glu and 10% ultralow IgG HI-FBS (Thermo Fisher Scientific, Waltham, MA), and pelleted at 350xg for 5min. The sample was resuspended in growth media and B cells were negatively selected using the EasySep Human B cell Isolation Kit according to the manufacturer’s protocol (STEMCELL Technologies, Vancouver, BC). Because our samples contained greater than one million PBMCs, B cell enrichment was used to provide a corresponding enrichment of B cell RNA in the total RNA added to the reverse transcription reaction. The final cells from negative enrichment were pelleted at 350xg for 5min, resuspended and lysed in Qiagen RLT buffer with beta-mercaptoethanol for 10min, frozen on dry ice, and transferred to −80°C storage until RNA purification with the Qiagen AllPrep RNA/DNA kit (Qiagen, Hilden, Germany). For generating stimulated PBMC samples, the day before PBMCs were thawed, 1.2 x10^6^ feeder cells were seeded in a T25 flask (VWR) in 4ml of B cell growth media and cultured overnight in a humidified 37°C, 5% CO_2_ incubator. PBMCs were quickly thawed at 37°C, washed 1x in 10ml of growth media, and resuspended in 4ml of 2x B cell growth media containing 2xITS from100x Insulin, Transferrin, Selenium (Thermo Fisher Scientific), 20ng/ml IL-10, 2ng/ml IL-2, 10ng/ml IL-15, 10ng/ml IL-6 (R&D Systems, Minneapolis, MN) and 4μg/ml CpG (ODN 2006-G5, InvivoGen, San Diego, CA). The 4ml of PBMCs were then added to the T25 flask with 4ml of conditioned feeder cell media. The final 8 ml cell culture was allowed to grow for 5 days at 37°C, 5% CO_2_ in a humidified incubator. At day 5 the cells were pelleted at 350xg for 5min, resuspended and lysed in Qiagen RLT buffer with beta-mercaptoethanol for 10min, frozen on dry ice, and transferred to −80°C storage until RNA purification. Since the culture conditions specifically expand memory B cells at the expense of viability of other cell types, no B cell enrichment was utilized at this step. Supernatants were also harvested to assay total immunoglobulin IgG, IgA and IgM secretion by indirect ELISA, as described below.

### Indirect ELISA Measurement of Total IgG, IgA and IgM

96-well, Nunc Maxisorp™ plates (VWR, Radnor, PA) were coated overnight with anti-human IgG Fc (Jackson ImmunoResearch, West Grove, PA), anti-human IgA Fc or anti-human IgM Fc (Bethyl Labs, Montgomery, TX) at 2μg/ml in PBS, pH7.2. The next day the plate was washed 3×300μl PBST and blocked for 1 hour in 1% BSA/PBS. A standard curve was prepared using human reference serum (Bethyl Labs) in 1/3 dilutions starting from targeted serum dilutions to give 200ng/ml of IgG, IgA or IgM, in assay diluent (0.5% BSA//PBS/0.05% Tween-20). Supernatants from stimulated PBMCs were diluted in assay diluent between 1/5 and 1/100. Both standards and samples were allowed to bind for 2 hours, washed 6x 300μl PBST and a 1/5000 cocktail of anti-human kappa-HRP and lambda-HRP antibodies (SouthernBiotech, Birmingham, AL) added for 1 hour in assay diluent. After 6x 300μl PBST washes, the plate was developed with KPL SureBlue™ TMB (VWR).

### BCR Primer Design and Pool Preparation

Dry oligos of desalted purity (IDT) were reconstituted at 100μM in Qiagen EB and stored in aliquots at −80°C. The oligos are shown in Table S1 and contain random base pairs to act as unique molecular identifiers (UMIs). The UMIs are added in variations of 8 or 12 nucleotide stretches to offset the high level of sequence similarity and lower Illumina sequencing accuracy in immunoglobulin amplicons at the 3’ and 5’ ends. A pool of IgH RT primers was made by mixing 10μl of each primer from the individual 100μM stocks (100μl final volume). Separately 10μl of each of lambda RT primers were mixed from individual 100μM stocks (20μl final volume). Next, a 10μM, 5:1 molar mix of Ig heavy:lambda chain RT primers was made using 16.7μl IgH RT primer pool and 3.3μl lambda RT primer pool, in a final 180μl of Qiagen EB. The same procedure and molar ratio were repeated in the preparation of the IgH (n=12): lambda (n=16) forward primers. Kappa RT and forward primer pools were prepared by mixing 10μl of each kappa RT (n=2) primer or kappa forward (n = 8) primer from the individual 100μM stocks and then diluting the mix to 10μM final.

### BCR Amplicon Preparation

Total RNA yields from the PBMC and stimulated PBMCs were in the range of 3 micrograms (50-100 ng/μl in 30μl Qiagen EB buffer) determined by absorbance at 260nm on the NanoDrop^TM^ One (Thermo Fisher Scientific). An input of 100-200ng total RNA was used for first strand cDNA synthesis with gene-specific reverse transcription (RT) primers directed to the constant regions. The RT primers for IgG, IgM, IgA and lambda were pooled, whereas the kappa RT was done in a separate reaction. Light chain kappa RT was carried out in separate reactions because the transcript abundance and amplification efficiency tended to out-compete heavy and lambda chains in multiplexed reactions. Primers are shown in Table S1. One to two hundred nanograms total RNA was added on ice to 10μM of pooled RT primers for HC/lambda or kappa chain (primer pools as described above and in Table S1) and 1mM of dNTP in a 10μl final volume, allowed to anneal for 3min at 72°C, and returned to ice. First strand reverse transcription was performed using SuperScript III RT (200U/μl, Thermo Fisher Scientific). To the 10μl annealed sample, on ice, 4μl of 5x Superscript RT buffer, 1μl 0.1M dithiothreitol, 1μl Superase-IN (20U/μl, Thermo Fisher), and 3μl of RNase free water were added, to give a final volume of 20μl. cDNA was made in a thermocycler for 1 hour at 50 °C, 5min 85°C, 4 °C hold. Second-strand cDNA was synthesized using Phusion High Fidelity Polymerase (Thermo Fisher). To the 20μl first strand cDNA,10μl of 5X Phusion buffer, 1μl of 10mM dNTPs, 0.5μl Phusion Taq, 1.5μl DMSO, 7μl of RNase free water, and 10μl of the 10μM pool of forward primers (as described above and shown in Table S1), were added, to give a final volume of 50μl. Samples were incubated at 98 °C for 4min, 52 °C for 1min, 72 °C for 5min and 4°C hold. Double-stranded cDNA was transferred to a low retention DNase-free 1.5ml Eppendorf tube and purified two times using Agencourt AMPure XP beads (Beckman Coulter, Brea, CA), at a volume ratio of 1:1, and eluted in 25μl of Qiagen EB buffer. Double-stranded cDNA was PCR amplified with Platinum DNA Polymerase High Fidelity (5U/μl HiFi Taq, Thermo Fisher). To the 25μl of eluted 2^nd^ strand cDNA, 5μl of 10x HiFi Taq buffer, 2μl of MgSO4, 1μl 10mM dNTPs, 0.2μl HiFi Taq, 1μl each of two PE primers completing Illumina adapter sequences (Table S1) and 14.8μl of water, were added, to give a final volume of 50μl. Samples were run at 94 °C for 2min, 27 cycles of 94°C for 30 s, 65 °C for 30s, and 68 °C for 2min, followed by 68 °C for 7min and 4°C hold. Final libraries were run on 2% E-Gel EX agarose gels (Thermo Fisher Scientific) and bands extracted with Quantum Prep Freeze and Squeeze DNA Extraction Spin Columns (BioRad, Hercules, CA). After one clean-up with 1:1 Agencourt AMPure XP beads, amplicons were eluted in 25-35μl Qiagen EB. An aliquot was diluted to 5-500pg/μl and 2μl quantified on the Agilent Fragment Analyzer Automated CE System using the DNF-474 High Sensitivity, 1bp-6000bp, NGS Fragment Analysis Kit (Advanced Analytical Technologies, Agilent Technologies, Santa Clara, CA), according to the manufacturer’s instructions. Pairs of samples with PBMC and stimulated PBMC amplicons were sequenced together using different Illumina barcodes to demultiplex after sequencing. Amplicon mixtures corresponding to 10:1 ratios of heavy chain + lambda: kappa were submitted for 300 forward × 250 reverse sequencing with MiSeq v3 kits (Illumina) at the Chan Zuckerberg Biohub Genomics Center. Addition of 15-20% PhiX was added to increase sequence diversity and overall sequencing performance. Each MiSeq run resulted in 7.5-20 million paired raw reads, which was reduced to 0.5-3.5 million unique immunoglobulin (Ig) sequences after processing.

### BCR Repertoire Data Analysis Pipeline

After completion of MiSeq sequencing, antibody repertoires were analyzed using methods based on the Immcantation pipeline (Vander Heiden *et al* 2014; Gupta *et al* 2015). An overview of BCR sequencing analysis and practical considerations, included in the Immcantation pipeline, are reviewed in Yaari G. and Kleinstein S.H. (2015). This pipeline continues to be updated as the field advances, and is composed of multiple software packages: pRESTO, Change-O, SHazaM, TIgGER, and Alakazam, found at https://immcantation.readthedocs.io. Because the Immcantation pipeline can be run using Docker containers, we created a cloud-based workflow incorporating Reflow (https://github.com/grailbio/reflow) that allowed for seamless processing of the constituent Immcantation software packages. The workflow is available at Github (https://github.com/czbiohub/bcell_pipeline). Some key characteristics of our workflow include the use of unique molecular identifiers (UMIs; Kinde *et al* 2011; Vollmers *et al* 2013) at both 5’ and 3’ ends of the Ig sequences, the collapse of sequences with identical UMIs, the use of the IgBLAST algorithm (Ye *et al* 2013) to calculate general Ig characteristics of each read, and the determination of clonal families by first calculating a clonal threshold distance via a nearest-neighbor algorithm and then collapsing sequences based on this threshold (Gupta *et al* 2017). We ran the initial steps using the pRESTO script (presto-abseq.sh at https://bitbucket.org/kleinstein/immcantation/src/97a70949607b6671a182a84d5052b705d1677891/pipelines/?at=default) with variations that are included in our Github repository. Given our sample and amplicon preparation included UMIs of varying lengths at both 5’ and 3’ ends, we included code to standardize the UMI length for subsequent steps (8bp at each end). The script first removes reads with average QC less than 20, and then annotates the reads based on 5’ and 3’ amplicon primers. All reads with identical UMIs are then collapsed with consensus sequences created and UMI numbers annotated into the sequence name. This is followed by assembly of paired-ends, at which point the UMIs at both ends are combined to create a 16bp signature per cDNA transcript, also annotated into the sequence name. Next the constant regions are re-checked for each paired read and isotype and subtype annotated into the sequence name. The final pRESTO steps include collapsing of identical sequences followed by a filter step to only include sequences found with 2 or more UMIs, to avoid including sequences that vary only due to sequencing error. The workflow continues with subsequent Immcantation packages, using the following scripts without changes at the website above: Change-O IgBLAST (changeo-igblast.sh), which calculates Ig repertoire characteristics, TigGER (tigger-genotype.R), which estimates novel V-gene alleles, SHazaM (shazam-threshold.R), which determines the optimal threshold for delineating clonal families, and Change-O Clone (changeo-clone.sh) that groups the sequences into clonal families. Lastly, we use a series of R scripts based on Alakazam that can be found at our Github page to visualize results. In addition to the usual Immcantation procedure of optimizing the clonal threshold value during each analysis (using SHazaM), we used a second strategy for comparisons across samples to identify mAb matches to clonal families in the repertoire. For these comparisons we included all unique heavy chain sequences (i.e. not only those with 2 or more UMIs), appended the mAb sequences, and applied a constant 12% threshold value to delineate clonal families. This was necessary because the optimized clonal thresholds applied in the primary analyses are not comparable across samples.

### Preparation of Samples for Bulk Transcriptome Analysis

Double stranded cDNA was prepared according to the single B cell cDNA synthesis described below. cDNA was diluted to range of 0.1-0.3 ng/μl, before tagmentation and PCR amplification with index primers using the Nextera XT DNA SMP Prep Kit and Nextera XT IDX Kit (Illumina, San Diego, CA). After cleanup with Agencourt AMPure XP beads (Beckman Coulter, Brea, CA) the sample was eluted in Qiagen EB buffer and quantified on the Agilent Fragment Analyzer Automated CE System using the DNF-474 High Sensitivity, 1bp-6000bp, NGS Fragment Analysis Kit (Advanced Analytical Technologies, Agilent Technologies, Santa Clara, CA), according to the manufacturer’s instructions. Libraries were submitted for 75×75bp sequencing on the Illumina NextSeq High Output instrument.

### Single Memory B cell Isolation

Human PBMCs were thawed and stained for 1 hour on ice in PBS, 1% BSA (w/v) with antibodies for positive markers of IgG memory B cells, and antibodies to negatively select and avoid contamination with dead cells, T cells, NK cells or monocytes (live, CD3^−^,CD14^−^,CD56^−^, IgM^−^, IgA^−^, CD19^+^,CD20^+^,CD27^+^). Biotinylated HA trimer (A/New Caledonia/20/1999) was complexed using 50nM at a 4:1 molar ratio with Streptavidin. Specific antibody clones and labels are listed in Table S2. The SH800 Cell Sorter was used with 100μm sorting chips (SONY Biotechnology, San Jose, CA) and NERL™ Diluent 2/Sheath Fluid for Flow Cytometry, (Thermo Fisher). Laser compensation was performed using the AbC™ Total Antibody Compensation Bead Kit (Thermo Fisher). Single cells were sorted directly into Hard-Shell® 96-well, low profile, thin wall, skirted, green/clear, PCR plates (BioRad) with 4μl of lysis buffer, frozen immediately on dry ice and stored at −80°C. cDNA was synthesized according to published Smart-seq2 methods (Picellli S. *et al* 2014). cDNA was cleaned up using 18μl of Agencourt AMPure XP beads (Beckman Coulter, Brea, CA) in the 25μl final cDNA volume, washed twice with 170μl 80% ethanol and eluted in 16μl of Qiagen EB. Heavy and light chain variable domains were amplified (95°C 5min, 30 cycles: 95°C 15s, 57°C 30s, 68°C 1min; and a final 10min 68°C extension) from 2-3μl of cDNA in 25μl of Accuprime Pfx supermix (Invitrogen, Thermo Fisher Scientific) and 0.5μl of the V_H_, V_kappa_ or V_lambda_ primer stock mixes described in Table S3. Sanger sequencing primers are also listed in Table S3.

### Recombinant Antibody Cloning, Production and Purification

Variable heavy or light chain domain PCR products were re-amplified in a nested PCR using primers with at least a 15 base pair overlap matching the 5’ signal sequence and 3’constant region of our human IgG_1_, kappa or lambda expression vectors (see Table S3 for primers). Expression vectors used were in-house constructs of Genbank LT615368.1, deposited by Tiller,T., *et al* (2009). Assembly of the gene fragments into expression vectors was performed with In-Fusion HD Cloning (Takara Bio USA, Mountain View, CA) and confirmed by Sanger sequencing (Sequetech, Mountain View, CA). Miniprep DNA (5μg) for the HC and LC of each mAb was transfected into 5ml of suspension Expi293 cells according to the manufacturer’s instructions (Thermo Fisher Scientific) in 50ml tubes. Cultures were grown in a Multitron shaker (INFORS HT, Annapolis Junction, MD) for five to eight days before supernatants were harvested, clarified, purified and assayed for HA binding. Antibodies were batch purified by adding 500μl of 1:1 slurry of MabSelect™ SuRe™ (GE, Chicago, IL) overnight. Supernatants and beads were decanted and rinsed into 10ml Poly-Prep ® Chromatography Columns (BioRad, Hercules, CA), washed with 20 volumes of PBS and eluted in 5ml of 20mM citrate buffer, pH3, neutralized with 250μl of 75mM 1.5M Tris-HCl, pH8.8. Purified antibodies were concentrated and exchanged by centrifugation into PBS, pH7.2 using Amicon Ultra-15 Centrifugal Filter Units with Ultracel-10 membranes (MilliporeSigma, Burlington, MA). IgG concentrations were determined by OD280 on the Nanodrop One.

### ELISA Measurement of mAb Binding to HA

96-well, Nunc Maxisorp™ plates were coated overnight with 1μg/ml H1N1 A/New Caledonia/20/1999 hemagglutinin (HA) trimer in PBS, pH7.2. HA was produced in-house by transiently transfecting Expi293 cells with the extracellular region of HA containing a C-terminal foldon domain for trimerization (Whittle J.R. *et al* 2014), Avi-tag for biotinylation and 6xHIS tag for Nickel purification using the HisPur™ Ni-NTA Spin Purification Kit (Thermo Fisher Scientific, Waltham, MA). The coated plates were washed 3x 300μl PBST and blocked for 1 hour in 1% BSA/PBS before adding a 1/3 titration of mAbs in assay diluent (0.5% BSA/PBS/0.05% Tween-20) for 1 hour at ambient temperature. Plates were washed 6x 300μl PBST and developed with secondary antibodies and TMB as described above. To measure the binding of mAbs in the presence of MEDI8852 Fab, the MEDI8852 IgG was cut to severe all Fc with LYSYL endopeptidase (Wako Chemicals, Richmond, VA) for 2 hours at 37°C. Plates were coated and blocked as described in the binding assay. Each mAb was prepared at a fixed concentration in 0.5% BSA/PBS/0.05% Tween-20 corresponding to a binding OD of 2. A 1/3 dilution series of MEDI8852 Fab was prepared starting from 100nM. Each mAb was added 1:1 to the Fab dilution series and allowed to bind for 1 hour at ambient temperature. Plates were washed 6x 300μl PBST and developed with an anti-human Fc specific secondary antibody (Invitrogen, Carlsbad, CA) and TMB as described above. Control wells for Fab binding alone were assayed alongside and the final signal subtracted from each corresponding dilution.

## RESULTS

### Experimental Rationale of PBMC BCR Repertoires +/− Stimulation

PBMCs from three normal donors of different genetic backgrounds 536 (African American male, 33yrs), 147 (Caucasian female, 21yrs.), and 682 (Hispanic male, 30yrs.) were purified and cryopreserved from Leukapheresis packs. Blood from donor 536 was collected at three different times, in November 2017, May 2018 and August 2018, while donors 147 and 682 were collected once, in August 2018. PBMC vials of 10^7^ cells were processed in one of four ways to obtain (i) PBMC B cell repertoires; (ii) stimulated PBMC B cell repertoires; (iii) bulk transcriptome profiles; or (iv) single, antigen-specific memory B cells. Two vials of cells were processed for each repertoire condition for each of the three donors, in separate experiments, to replicate data. Examples of typical BCR amplicon products used for NGS, which looked similar between PBMC and stimulated PBMC, are shown in Figure 1. The analysis of immunoglobulin repertoires from NGS data is described in detail in the Methods section and a summary of the workflow is provided in Figure 2. Our PMBC stimulation was based on ELISPOT methods, previously demonstrated to give rise to the selective expansion, identification and cloning of antigen-specific memory B cells (Crotty S. *et al* 2004, Weiss G.E. *et al* 2012, McCutcheon K.M. *et al* 2014). Total IgG, IgM and IgA secreted by activated B cells was measured between 82-954, 29-266, and 8-35 ng/ml among replicate stimulated cultures of the three donors, respectively (Table 1). The functional *in vitro* activation of the PBMCs was further characterized by mRNA expression. Total RNA was used to compare the bulk cellular transcriptome in one replicate of each of the three donors PBMCs before and after stimulation. Upon stimulation, transcripts corresponding to both innate and adaptive immunity were upregulated 10-1000-fold over unstimulated PBMCs. Representative genes for T cell and T cell Receptor (TCR) activation, BCR activation and B cell differentiation, as well as genes involved in various innate immune cell activities (e.g. Natural killer cells, Dendritic cells, and Granulocytes) are shown in Table 2. The stimulation of PBMCs to increase representation of memory B cells should correspond to an improved ability to identify monoclonal antibody sequences in NGS repertoires. To test this, eight antigen-specific mAbs to the hemagglutinin trimer of influenza H1N1 were obtained by FACS from one donor (536, November blood draw sample), the heavy and light chains sequenced from single cells, and recombinant mAbs made. Their binding activity was confirmed by ELISA. The mAbs, derived from several different IGVH germlines, demonstrated ELISA binding EC_50_ values in the range of 5-20nM, comparable to the high affinity, highly cross-reactive, MEDI8852 mAb directed to the HA stalk region (Figure 3A). MEDI8852 was made in-house using the published sequence (Kallewaard N.L 2016) and is shown with black dotted lines in Figure 3. The binding of antibodies INF3, INF7, INF9 and INF11 to the HA trimer demonstrated blocking by the Fab fragment of MEDI8852 (Fab blocking the intact MEDI8852 mAb is shown in black), indicative of common binding to the HA stalk region (Figure 3B). At high Fab concentrations most mAbs reached a plateau level of binding, including the IgG form of MEDI8852. This likely reflects a steady state level of Fab displacement by IgG binding with higher affinity and avidity during the one hour incubation time. Alternately, some mAbs may be able to retain a low level of equilibrium binding to partial epitopes outside the stalk region. Data identifying these memory B cell mAbs in the repertoires of donor 536 is reported below.

**TABLE 1.** Immunoglobulin in Leukopak Supernatant and Secreted from Stimulated PBMC Cultures.

|  | Leukopak supernatant ug/mL |  |  |  |  | 5-day B cell stimulated media ng/mL |  |  |  |  |
| --- | --- | --- | --- | --- | --- | --- | --- | --- | --- | --- |
|  | 536N | 536M | 536A | 147A | 682A | 536N | 536M | 536A | 147A | 682A |
| IgG | 192 | 129 | 121 | 138 | 190 | 448 | 329 | 278 | 185 | 417 |
| (replicate) | - | - | - | - | - | 954 | 374 | 82 | 123 | 362 |
| IgM | 483 | 296 | 273 | 212 | 240 | 163 | 96 | 57 | 29 | 57 |
| (replicate) | - | - | - | - | - | 266 | 97 | 45 | 34 | 55 |
| IgA | 22 | 11 | 14 | 15 | 25 | 24 | 19 | 13 | 8 | 13 |
| (replicate) | - | - | - | - | - | 35 | 14 | 11 | 9 | 11 |

*N= November; M= May; A= August*

**TABLE 2.** RNAseq Transcript Reads in Innate and Adaptive Immune Responses between PBMCs and *In Vitro* Stimulated PBMCs.^1^.

| GENE | donor 147 |  | donor 536 |  | donor 682 |  |
| --- | --- | --- | --- | --- | --- | --- |
|  | PBMC | stim-PBMC | PBMC | stim-PBMC | PBMC | stim-PBMC |
| **Immune Cells** |  |  |  |  |  |  |
| C1QC | 2 | 1113 | 1 | 121 | 2 | 132 |
| CCL2 | 6 | 3912 | 7 | 1254 | 24 | 911 |
| CCR2 | 11 | 222 | 13 | 137 | 18 | 292 |
| CCR5 | 7 | 245 | 1 | 215 | 12 | 118 |
| CD209 | 19 | 661 | 8 | 141 | 6 | 111 |
| CD96 | 75 | 1927 | 195 | 1885 | 163 | 834 |
| CXCL1 | 1 | 839 | 1 | 512 | 4 | 148 |
| CXCL8 | 28 | 5419 | 3 | 4821 | 1448 | 517 |
| FCGR1A | 2 | 115 | 1 | 22 | 11 | 25 |
| GNLY | 18 | 6616 | 12 | 11662 | 293 | 4512 |
| GZMB | 6 | 4569 | 5 | 9192 | 41 | 4494 |
| IL7R | 26 | 2128 | 31 | 783 | 236 | 1291 |
| LAG3 | 4 | 292 | 9 | 296 | 4 | 274 |
| LEF1 | 19 | 1676 | 26 | 716 | 143 | 633 |
| PRF1 | 6 | 4242 | 14 | 5633 | 75 | 1921 |
| RSAD2 | 18 | 324 | 9 | 176 | 9 | 61 |
| SLA2 | 14 | 933 | 18 | 854 | 149 | 467 |
| SLAMF7 | 31 | 1216 | 76 | 1331 | 115 | 412 |
| XBP1 | 212 | 5119 | 531 | 8839 | 1319 | 7383 |
| XCL2 | 4 | 65 | 5 | 115 | 2 | 51 |
| ZAP70 | 38 | 2185 | 26 | 1613 | 312 | 1156 |
| **T cells** |  |  |  |  |  |  |
| CD247 | 5 | 321 | 6 | 1152 | 28 | 184 |
| CD274 | 7 | 3148 | 15 | 201 | 194 | 1205 |
| CD3D | 5 | 1318 | 5 | 628 | 131 | 647 |
| CD3E | 10 | 1250 | 4 | 988 | 182 | 585 |
| CD3G | 5 | 3770 | 14 | 440 | 41 | 2049 |
| CD7 | 5 | 3069 | 4 | 482 | 60 | 960 |
| CD8A | 7 | 809 | 8 | 2137 | 159 | 619 |
| CD8B | 4 | 591 | 11 | 1604 | 63 | 461 |
| CISH | 14 | 986 | 16 | 2497 | 111 | 456 |
| EOMES | 13 | 1746 | 9 | 517 | 177 | 759 |
| HAVCR2 | 30 | 1956 | 9 | 1006 | 93 | 1015 |
| LAT | 7 | 622 | 11 | 1213 | 33 | 245 |
| PRKCQ | 28 | 845 | 21 | 698 | 66 | 417 |
| SIT1 | 10 | 163 | 16 | 327 | 8 | 108 |
| SPN | 18 | 509 | 26 | 2025 | 47 | 289 |
| TRAT1 | 14 | 591 | 1 | 278 | 26 | 202 |
| TXK | 7 | 398 | 15 | 372 | 21 | 320 |
| ZNF683 | 17 | 438 | 6 | 195 | 19 | 123 |
| **B cells** |  |  |  |  |  |  |
| CD27 | 31 | 975 | 62 | 953 | 49 | 730 |
| CD38 | 16 | 602 | 49 | 801 | 11 | 358 |
| IGLL5 | 380 | 4817 | 610 | 10902 | 339 | 10939 |
| JCHAIN | 94 | 1506 | 283 | 5180 | 126 | 5065 |
| PRDM1 | 38 | 769 | 90 | 1280 | 169 | 710 |
| TNFRSF17 | 1 | 29 | 2 | 40 | 0 | 51 |
| UNG | 8 | 358 | 6 | 320 | 6 | 196 |

<sup>1</sup>August timepoint; Not normalized; Most highly upregulated genes from Adaptive Immune Response GO\_0002250.

**FIGURE 1.**
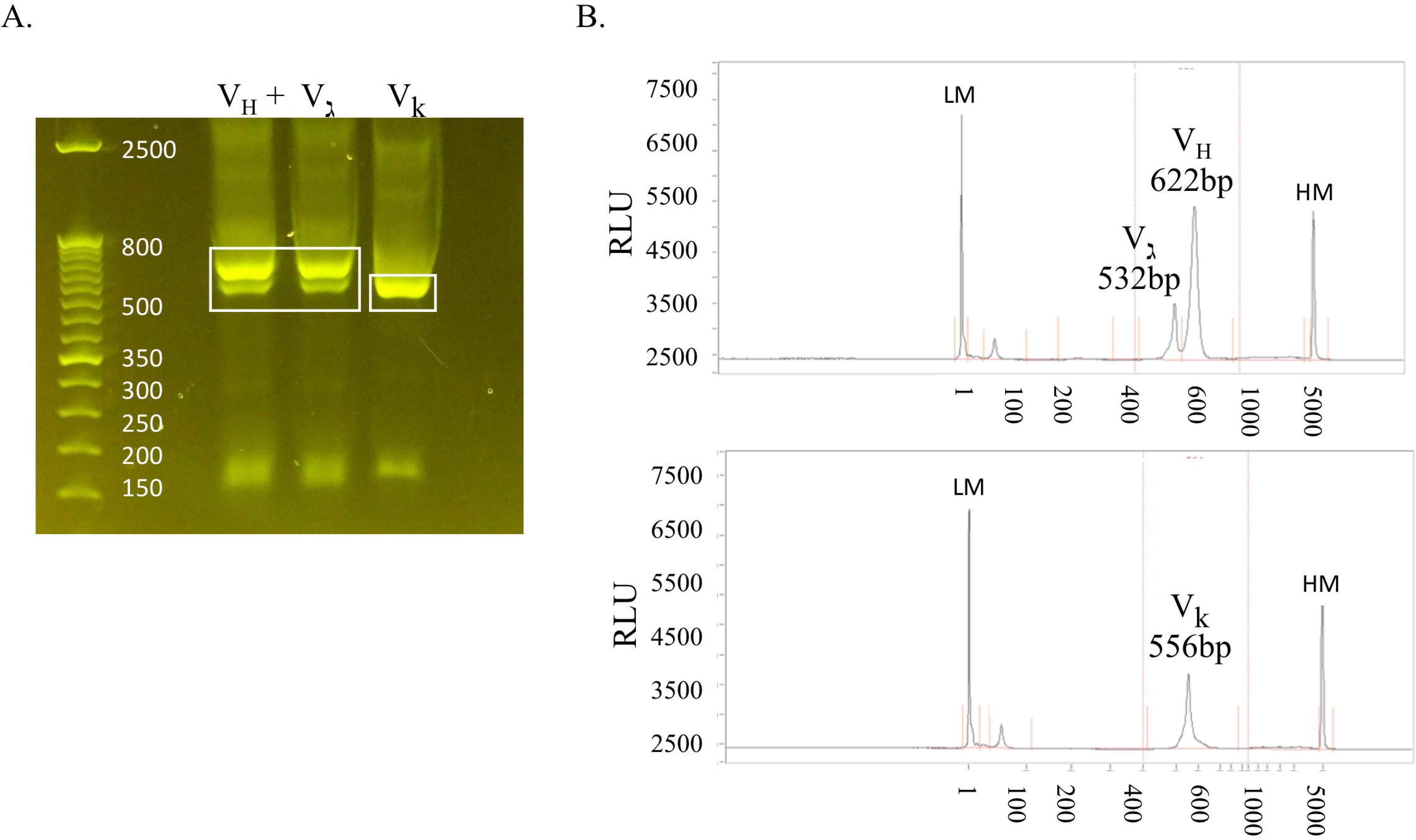
Amplicon agarose gel purification and fragment analyzer quantitation. A.

PCR amplicons of the human immunoglobulin heavy/lambda light chain and kappa light chain from total cellular RNA are shown on a 2% agarose gel. The white boxes indicate the regions excised from the gel for extraction. **B.** Examples of fragment analyzer results for the final extracted and purified amplicons immediately prior to sequencing submission are shown with the V_H_ at 622bp, V_lambda_ 532bp (upper plot) and V_kappa_ at 556bp (lower plot). Peaks labeled LM and HM are low and high molecular size markers, respectively. Areas under the curves are used for quantitation, with typical yields being in the range of 5ng/μl in 25-35μl.

**FIGURE 2.**
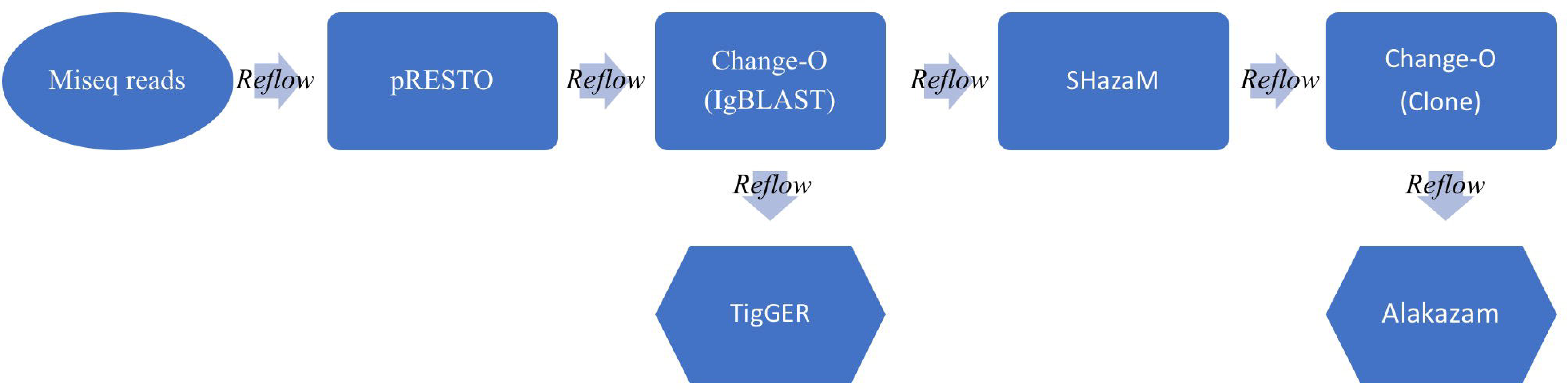
Schematic of the BCR analysis workflow.

Raw paired MiSeq reads (300 × 250bp) are inputted into the Immcantation pipeline. The pRESTO package processes the MiSeq reads and does QC, UMI processing, primer and isotype/subtype annotation & paired read assembly to output annotated sequence lists (.fastq). Change-O next runs the IgBLAST algorithm on all reads, annotating germline information to output tabular files (.tab). TigGER calculates novel germlines, while SHazaM calculates the optimal threshold for clonal clustering. Change-O then clusters the reads into clones and Alakazam summarizes the outputs in graphical form (.png &.pdf). Throughout the workflow, a Reflow script pieces all of the parts together, delineating outputs and inputs and allowing the pipeline to run in the cloud (using Amazon Web Services).

**FIGURE 3.**
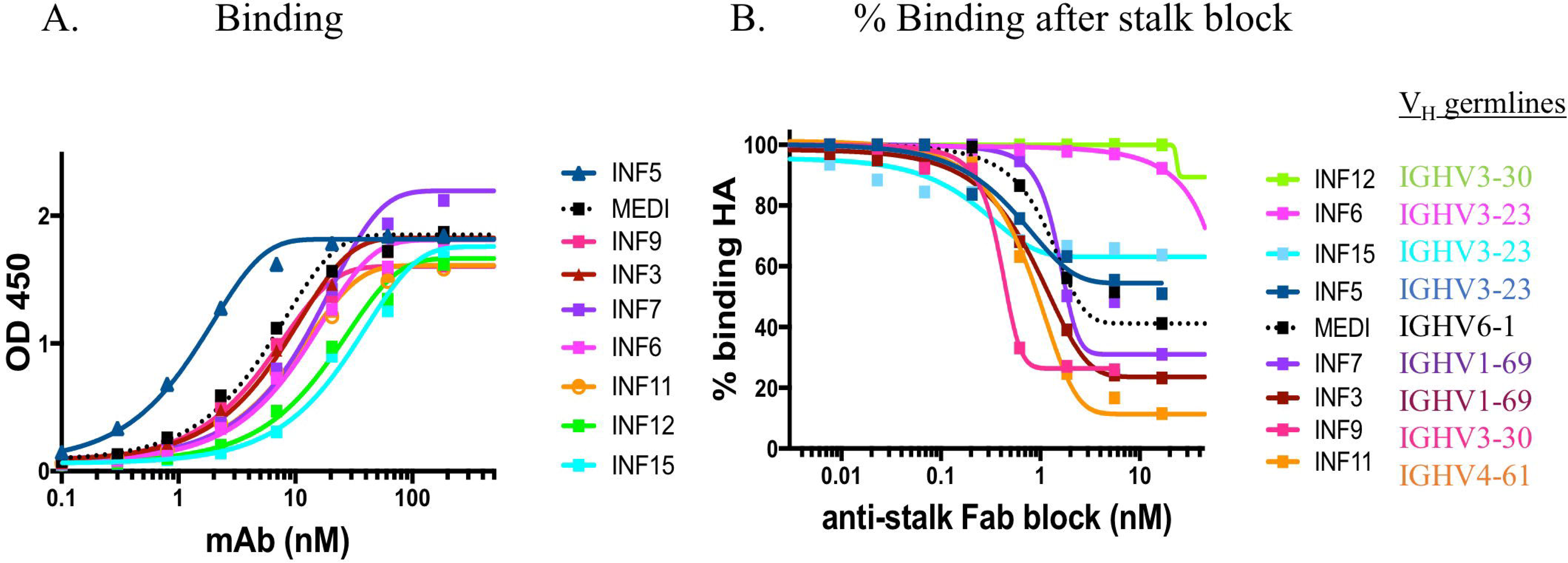
ELISA binding of mAbs to H1N1 HA and functional epitope mapping to the stalk region. A.

Purified recombinant monoclonal antibodies were tested for binding to the H1N1 HA trimer compared to the reference in house synthesized MEDI8852 IgG (shown in dotted black line). **B.** The binding of monoclonal antibodies was measured after blocking the coated HA trimer with increasing concentrations of MEDI8852 Fab. The binding of antibodies INF3, INF7, INF9 and INF11 to the HA trimer were effectively blocked by the Fab fragment of MEDI8852 (Fab blocking the intact MEDI8852 mAb itself is shown in the dotted black line), indicative of common binding of these mAbs to the HA stalk region. In this assay, only INF11 was close to completely blocked at high Fab concentrations. During the one-hour incubation time, higher avidity and affinity IgG (including MEDI8852 IgG) may have reached an equilibrium by displacing bound Fab or by binding to partial epitopes on the trimer independent of the stalk region.

### Equivalent Germline Usage in PBMC and Stimulated PBMC Repertoires

Comparing V-gene usage between repertoires, we found that in both PBMC and stimulated PBMC repertoires gene usage was consistent with previously described healthy BCR repertoire data (e.g. Briney B. *et al* 2019), and germline usage was equivalent across PBMC and stimulated PBMC repertoires (Figure 4). For example, examining IgG and IgM sequences in one replicate at one timepoint, VH1-2, VH3-23, VH3-33, VH4-59 and VH5-51 are among the most commonly used V-genes across both PBMC and stimulated PBMC repertoires in donor 147, VH1-2, VH1-69, VH3-30, VH4-39 and VH5-51 are among the most commonly used V-genes across both PBMC and stimulated PBMC repertoires in donor 536, and VH1-69, VH3-23, VH4-39, VH4-59 and VH5-51 the most commonly used V-genes across both PBMC and stimulated PBMC repertoires in donor 682 (Figure 4).

**FIGURE 4.**
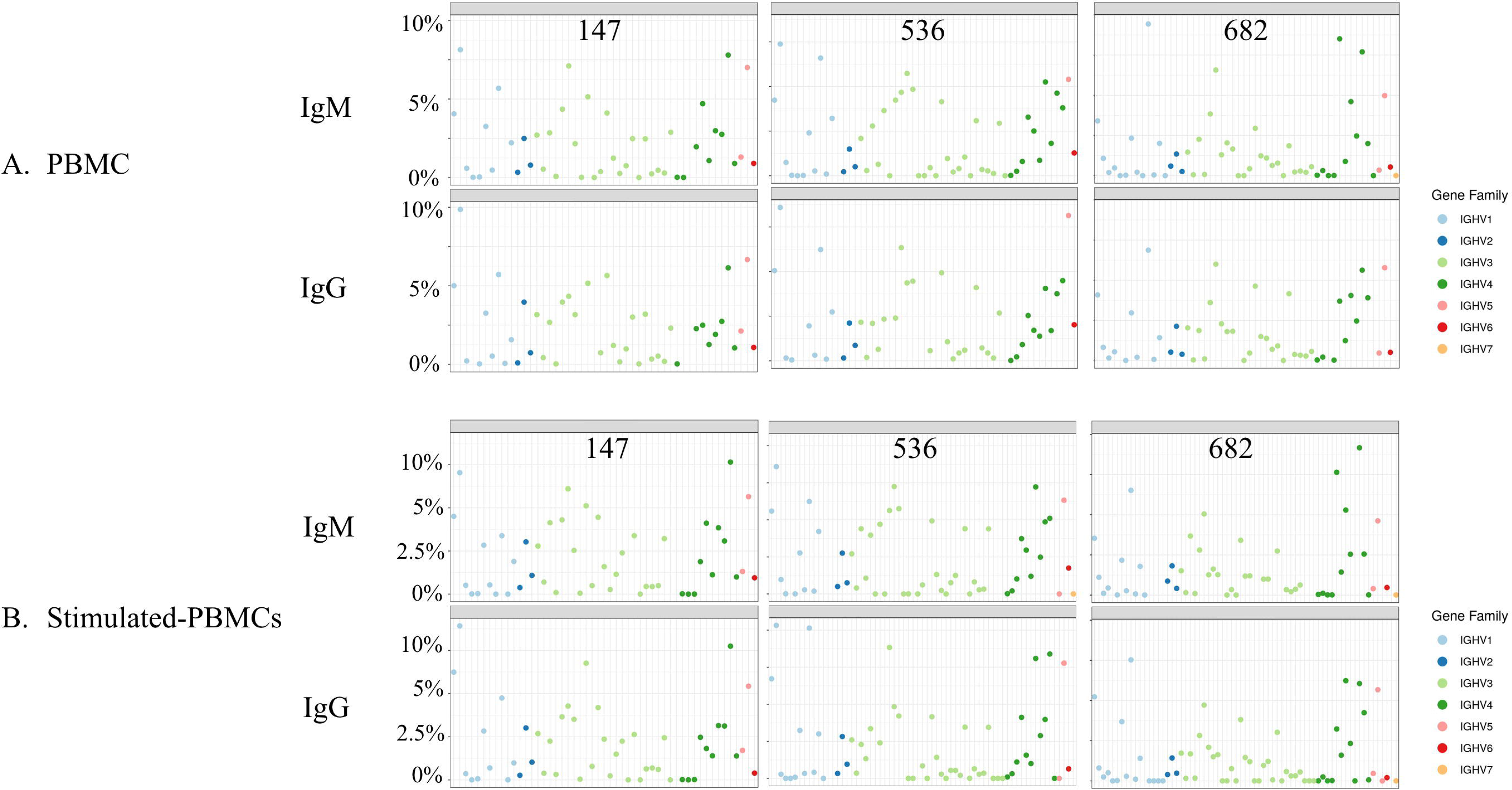
Germline V-gene usage among A. PBMC and B.

Stimulated PBMC repertoires. Plots show V-gene usage for one replicate of each of the three donor repertoires (147, 536 and 682) sampled at the August 2018 timepoint, and are separated by isotype, with only IgM and IgG reads shown. Plots were made using the Alakazam R scripts in the Immcantation pipeline.

### Stimulated PBMC Repertoires Shift to More Antigen-Experienced Clones

We examined whether PBMC and stimulated PBMC repertoires vary in their antigen experience by quantifying heavy chain isotype ratios and SHM among IgM and IgG sequences. Among three donors, we found a consistent shift in repertoires with respect to heavy chain isotype and subtype. In PBMC repertoires, IgM made up over 90% of all heavy chain reads across all samples and replicates. In contrast, stimulated PBMCs are greatly enriched for IgG (and particularly the IgG_1_ subtype), comprising between 58% and 82% of all heavy chain reads (Figure 5). Somatic hypermutation frequencies also significantly increased (p < 0.001 for all comparisons) in the 3 different donors, although to varying degrees, with PBMC repertoires averaging 0.6% SHM in IgM reads and 5.0% SHM in IgG reads but stimulated PBMC repertoires showing an increase in somatic hypermutation, averaging 2.0% SHM in IgM reads and 5.7% SHM in IgG reads (Figure 6). In our searches for single cell memory mAbs in donor 536, we found matching clonal families for the influenza mAbs most frequently among stimulated PBMC repertoires across all blood draw timepoints (Figure 7a). These hit rates improved despite the fact that the total number of clonal families in stimulated PBMC repertoires were 2 to 12-fold lower than PBMC (e.g. 204,000 vs 16,000 in a single August replicate for donor 536; Table 3), but were enriched for the IgG switched and higher SHM phenotype of antigen-experienced BCR (Figures 5 and 6). As would be expected from analyzing samples from a dynamic immune system, we observed differences in mAb matches to Ig repertoires among the timepoints. As shown in Figure 7a, mAb searches at the initial timepoint (November), when the mAbs were cloned, showed equal hits for PBMC and stimulated PBMC (4/8 mAbs). The second timepoint (May) had the greatest number of matches in the stimulated PBMC repertoire, (7/8 mAbs) relative to the PBMC (2/8), while the number at the third timepoint (August) dropped to 0/8 mAbs in PBMC and 2/8 mAbs in stimulated PBMC. Further analyses of the relative size and diversity of the mAb clonal families at the May timepoint, are described in sections below. The donor was collected as healthy and no medical history is known, but these findings could be consistent with this donor having received exposure or vaccination to influenza A virus just prior to the November blood draw, with the immune response waning in the PBMC compartment by the following August. In addition to identifying mAb in the stimulated PBMC repertoires more frequently, we also found considerably more reads matching these clonal families (Figure 7b). Out of 5 million total processed Ig reads (i.e. after the Change-O step shown in Figure 2) across PBMC repertoires for November, May and August, reads matching the mAb clonal families ranged from 0-18, while in stimulated PBMC repertoires, matching mAb reads ranged from 95 to 448 out of a total of 7 million reads. Activated B cells proliferate and significantly upregulate mRNA transcription, thus making clonal families of interest more easily found, and found in larger numbers.

**TABLE 3.** Summary of Ig sequences, clonal families and their corresponding mean SHM in PBMCs and *In Vitro* Stimulated PBMCs.

| **PBMCs** | **147A** |  | **536N** |  | **536M** |  | **536A** |  | **682A** |  |
| --- | --- | --- | --- | --- | --- | --- | --- | --- | --- | --- |
| HC reads | 90398 | 59651 | 112128 | 144519 | 139389 | 186664 | 253586 | 173944 | 239491 | 145578 |
| HC clonal families | 75672 | 49595 | 95095 | 102093 | 119668 | 148994 | 204129 | 145749 | 202146 | 129432 |
| IgM reads | 82607 | 53839 | 103577 | 132636 | 129498 | 174385 | 238019 | 163658 | 217268 | 131032 |
| SHM, IgM reads | 0.8% | 0.8% | 0.8% | 1.0% | 0.8% | 0.8% | 0.6% | 0.6% | 0.6% | 0.5% |
| IgG reads | 4768 | 3255 | 4760 | 7174 | 5481 | 7135 | 8735 | 6389 | 12916 | 7566 |
| SHM, IgG reads | 4.4% | 4.3% | 5.5% | 5.4% | 5.5% | 5.4% | 5.4% | 5.4% | 5.1% | 5.1% |
| IgM clones | 68789 | 44406 | 87385 | 92265 | 110789 | 138560 | 190087 | 136855 | 183286 | 117115 |
| SHM, IgM clones | 0.6% | 0.6% | 0.6% | 0.6% | 0.6% | 0.5% | 0.4% | 0.4% | 0.5% | 0.5% |
| IgG clones | 4231 | 2935 | 4277 | 5815 | 4900 | 6000 | 7286 | 5427 | 11505 | 6637 |
| SHM, IgG clones | 4.3% | 4.2% | 5.3% | 5.1% | 5.5% | 5.2% | 5.3% | 5.3% | 5.1% | 5.0% |
| **Stimulated PBMCs** | **147A** |  | **536N** |  | **536M** |  | **536A** |  | **682A** |  |
| HC reads | 128312 | 99898 | 71594 | 124013 | 185986 | 179813 | 51720 | 142435 | 58893 | 89055 |
| HC clonal families | 72805 | 41404 | 12626 | 19424 | 64264 | 86379 | 15838 | 51756 | 21798 | 26944 |
| IgM reads | 69335 | 34993 | 13295 | 28893 | 64401 | 85831 | 6039 | 54147 | 2430 | 13437 |
| SHM, IgM reads | 1.2% | 1.1% | 4.2% | 2.8% | 1.8% | 1.3% | 4.0% | 1.2% | 3.4% | 1.4% |
| IgG reads | 55161 | 62984 | 55293 | 89078 | 113665 | 87748 | 44655 | 84324 | 56846 | 73874 |
| SHM, IgG reads | 4.7% | 4.5% | 6.4% | 6.5% | 6.3% | 6.1% | 6.6% | 6.7% | 5.8% | 6.2% |
| IgM clones | 52584 | 27928 | 3267 | 12249 | 42532 | 63103 | 2034 | 38535 | 1258 | 10068 |
| SHM, IgM clones | 0.7% | 0.7% | 1.5% | 0.8% | 0.7% | 0.6% | 1.9% | 0.5% | 2.1% | 0.7% |
| IgG clones | 17331 | 12179 | 8557 | 5896 | 18706 | 19822 | 13321 | 11631 | 20438 | 15465 |
| SHM, IgG clones | 4.6% | 4.6% | 6.0% | 6.1% | 6.2% | 6.1% | 6.0% | 6.5% | 5.5% | 5.8% |

*N= November; M= May; A= August*

**FIGURE 5.**
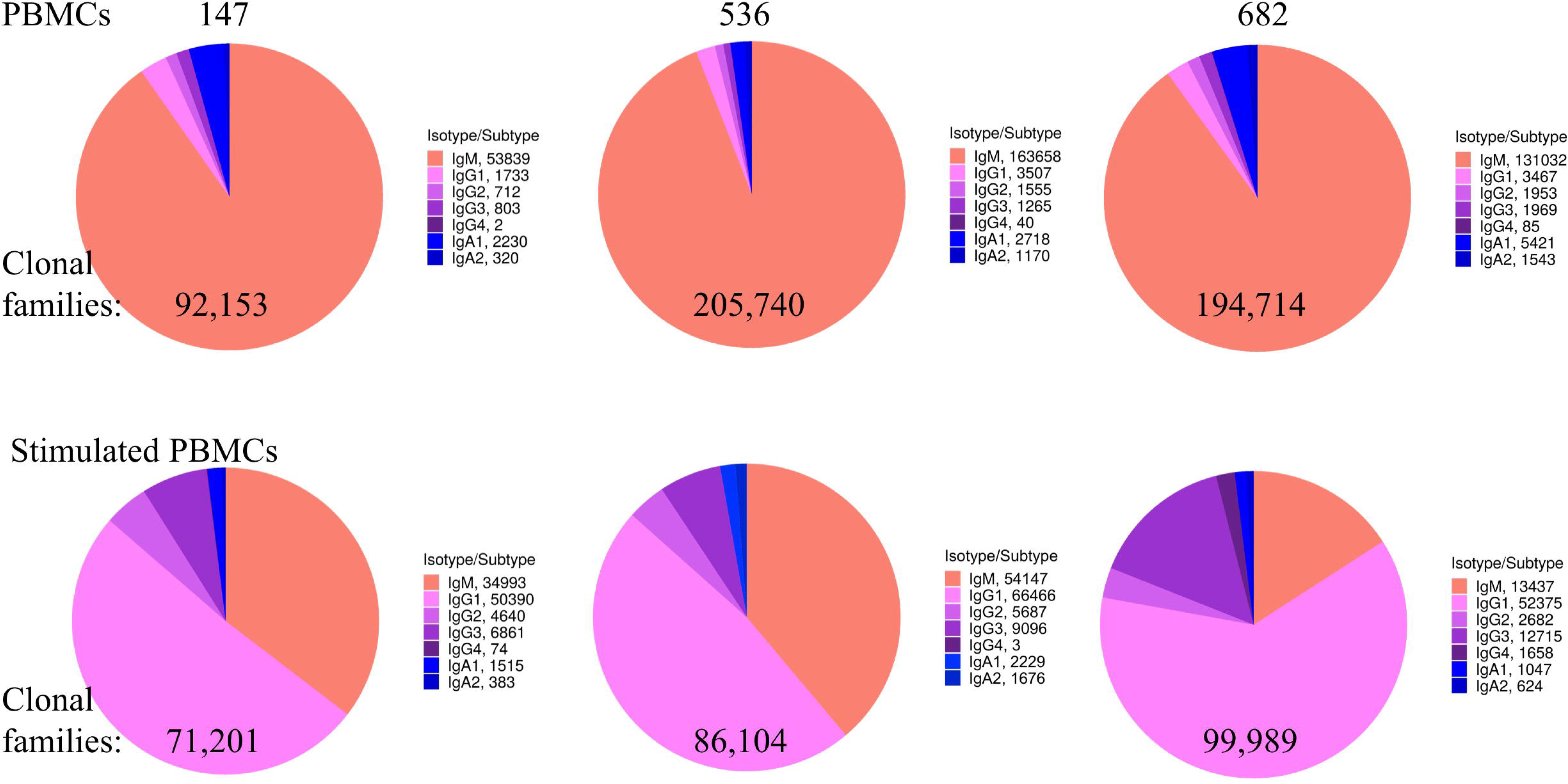
Heavy chain isotype and subtype usage among PBMC and stimulated PBMC repertoires.

Plots show the proportion of heavy chain isotypes for one replicate of each of the three donors (147, 536 and 682) sampled at the August 2018 timepoint. While pie chart numbers show unique reads found with >2 UMIs, the more stringent criteria for enumerating clonal families by V-D-J sequence similarity is also shown (in bold numbers on pie charts). A constant clonal threshold value of 12% was used to assign clones into families, so data could be compared across samples (see Methods). Plots were made using the Alakazam R scripts in the Immcantation pipeline.

**FIGURE 6.**
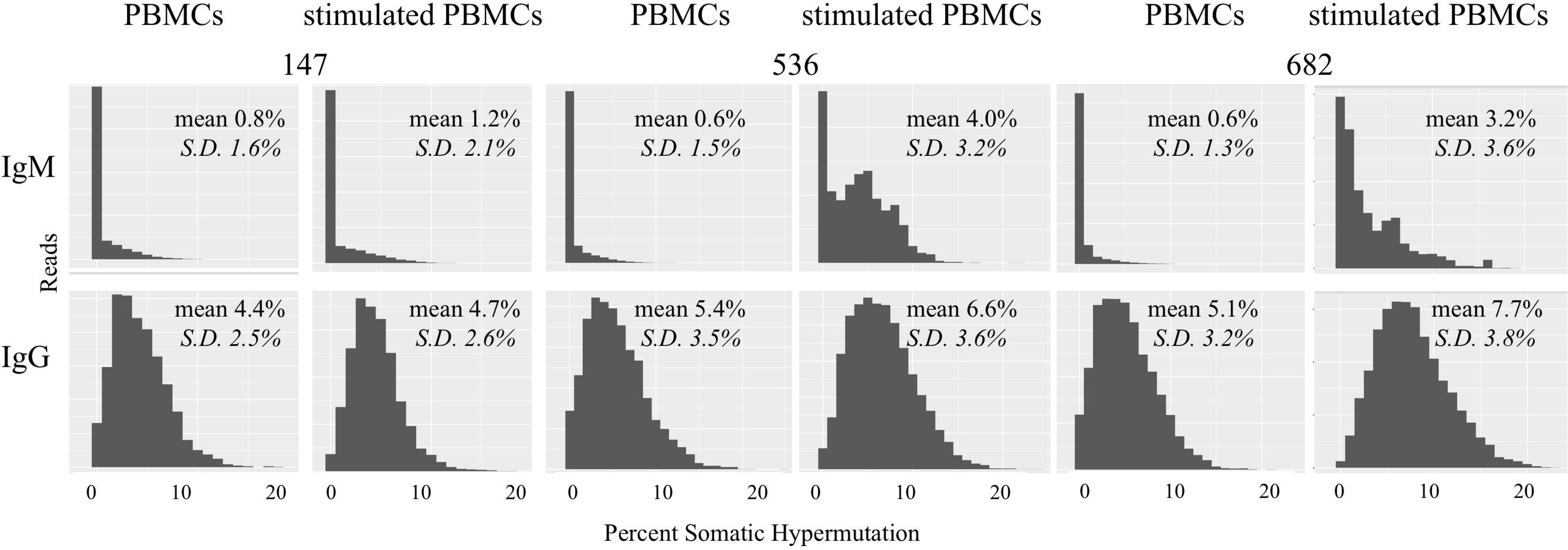
Somatic hypermutation among PBMC and stimulated PBMC repertoires.

Plots show distributions of SHM values across all IgM and IgG reads in PBMC repertoires vs. stimulated PBMC repertoires in one replicate, for each of the three donors (147, 536 and 682) sampled at the August 2018 timepoint. Plots were made using the Alakazam R scripts in the Immcantation pipeline. Since the total number of reads in each distribution plot are different, to compare across samples, the y-axes were normalized to the maximum number of reads.

**FIGURE 7.**
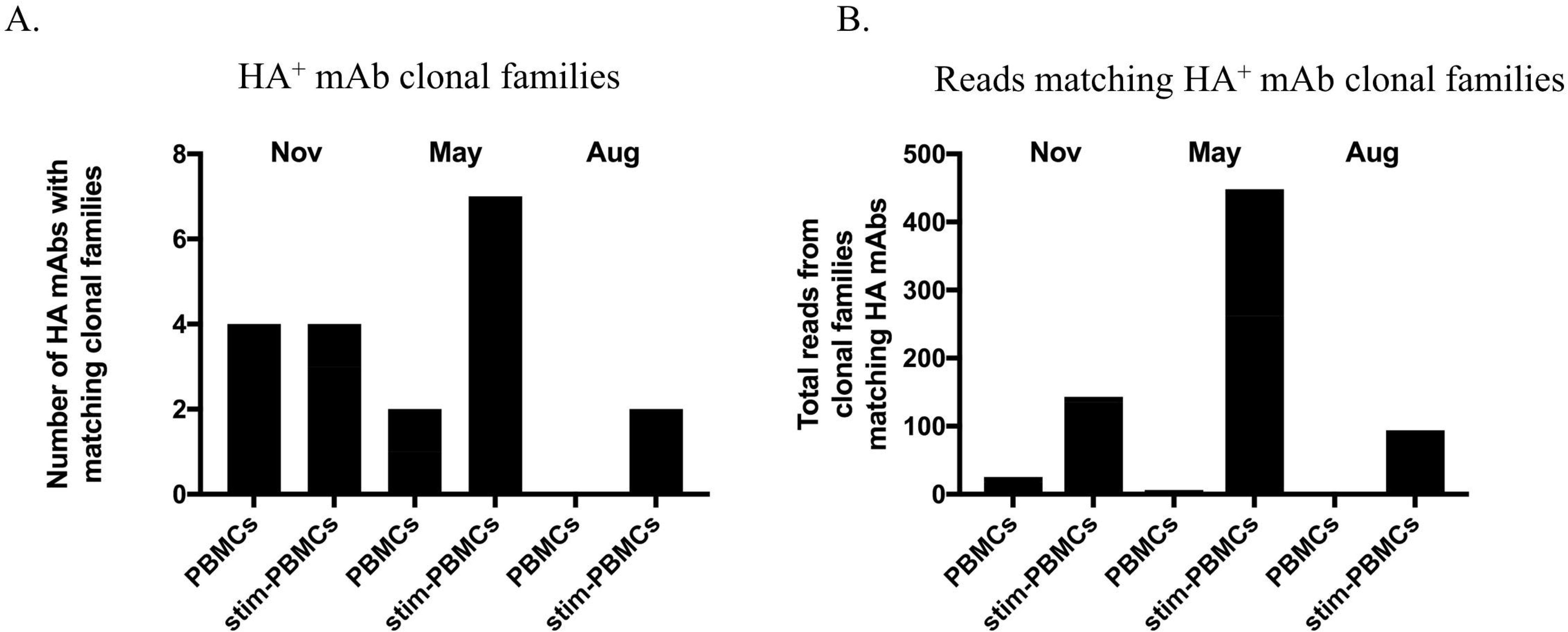
Clonal families of 8 HA^+^ memory B cell sequences found among PBMC and stimulated PBMC repertoires. A.

Number of mAbs out of the eight HA^+^ memory B cell sequences found with clonal families in either PBMC repertoires or stimulated PBMC repertoires, for all three timepoints of donor 536. One mAb (INF15) was not found in any of the repertoires. **B.** Number of reads found within these clonal families, in either PBMC repertoires or stimulated PBMC repertoires for all three timepoints of donor 536.

### Persistent B Cell Clones are Found More Frequently in Stimulated PBMC

With the three samples of PBMCs drawn from a single donor, we examined whether there were ‘persistent’ BCR sequences found across all timepoints spanning a 9 month period. Identical BCR sequences found consistently over time would more likely be derived from the memory B cell population, particularly if they have switched to the IgG isotype and have V-D-J sequences mutated from germline. Searches for identical BCR were conducted in Geneious v11.1 (Biomatters, Auckland, New Zealand) on data merged from the replicate experiments, using DNA segments corresponding to FR1 residue 12 to FR4 residue 136 (Kabat numbering). Among 5 million processed Ig reads from donor 536 PBMC across November, May and August, we found 7229 identical BCR sequences that persisted across all timepoints, of which only a small fraction (173) were mutated IgG BCR sequences. In contrast, out of 7 million corresponding donor 536 stimulated PBMC sequences, 2908 identical BCR sequences persisted across the three timepoints, of which the majority (2151), were mutated IgG BCR sequences (Figure 8). The BCR sequences had unique UMI usage between datasets, making it unlikely they were the result of cross-contamination. UMI codes are in the primers at the RT step where they are uniquely incorporated into a single cDNA. Also, the subsequent PCR reaction adds index primers to the UMI to differentiate reads between samples, further preventing incorrect sequence assignments downstream. It would be expected that, over this individual’s lifetime, the sequences of many of these persistent clones, observed herein over a 9 month period, would change. Longitudinal studies would be useful to evaluate lifetime immune repertoires circulating in PBMC, and how these relate to immunoglobulin sequences in serum (Wine Y. *et al* 2015, Boutz, D.R. *et al* 2014).

**FIGURE 8.**
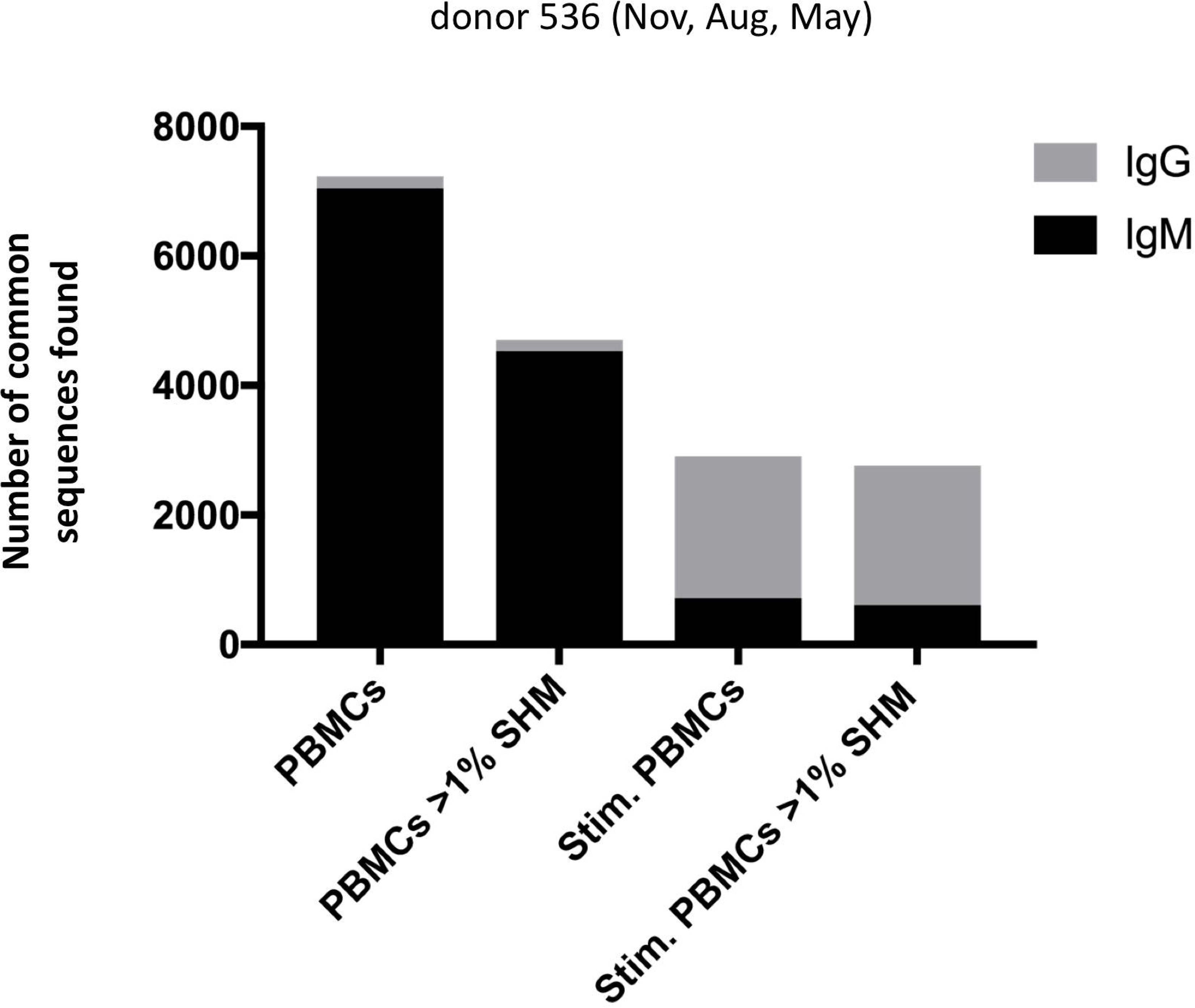
Number of persistent sequences found in all three timepoints in a single donor, among PBMC or stimulated PBMC repertoires.

Each column indicates the number of identical heavy chain sequences for IgM (black) or IgG (gray) clones found across all three timepoints of donor 536, for PBMC or stimulated PBMC. Either total reads or mutated reads (greater than 1% SHM) are shown as indicated on the x-axis.

### Stimulated PBMC Repertoire Data can be used to Determine the Immunological Hierarchy of B Cell Clonal Memory

The deeper memory BCR repertoire data obtained from stimulated PBMCs allows for the identification of large clonal families related to known antigen-specific mAbs, making it possible to examine the immunological context of antigen-specific mAbs in detail. For instance, we were able to map HA-specific mAbs onto lineages or networks of clonal families, highlighting aspects such as isotype, subtype and abundance (by UMI read counts). For example, the clonal lineage of HA stalk-region binding antibody INF9 is shown in Figure 9. The representation was made using the Alakazam package in the Immcantation pipeline and is a parsimony-based network of the reads clonally related to INF9 (in donor 536, single replicate of May timepoint), using data that could only be found in the stimulated PBMC repertoire. Within the lineage, INF9 itself appears as a minor player, with other clones having been selected for larger size and SHM divergence. Although clones of IgM and IgA isotype were observed, IgG dominated the response. We can also examine the relative abundances of antigen-specific mAbs in stimulated PBMC repertoires. Again, in a stimulated May repertoire of donor 536, containing 1 million reads and a total of 127,448 clonal families, we looked for reads related to the 8 antigen-specific mAbs (refer to Methods for mAb matching strategy). Heavy chain clonotypes were found for 7/8 mAbs, and plotted by both average SHM and by how many times they were found in the repertoire (Figure 10). In this example, mAb INF9 has the most abundantly expressed clonal family of the 8 antigen-specific mAbs, containing 166 different reads, and a 6% average SHM. In comparison, mAb INF12 shows the highest average SHM (12%) and low abundance (1 read of 1 clone) in the May repertoire. Despite being of low abundance in the May stimulated PBMC repertoire, divergent reads of variable UMI counts related to INF12 were found across all timepoints, indicating its persistence (data not shown). Thus, from these data it is possible to assess the relative immunodominant hierarchy of B cell clonal memory, which could be correlated to past and future protection from disease and better inform vaccine design.

**FIGURE 9.**
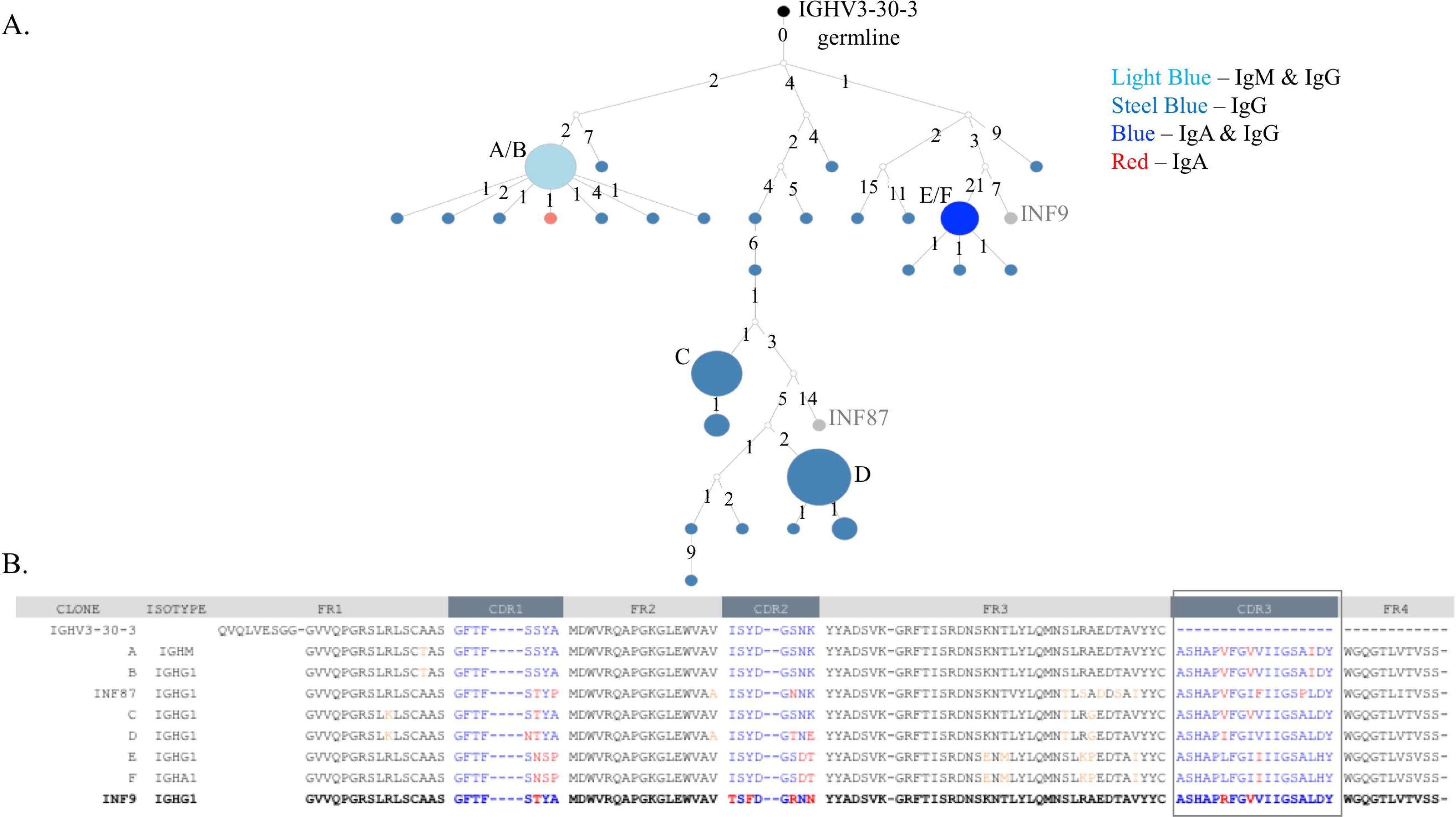
Clonal family members of HA^+^ memory B cell mAb INF9 found in a stimulated PBMC repertoire. A.

The clonal lineage of mAb INF9 deriving from the germline (IGHV3-30-3) in black, with IgG sequences in steel blue, IgM/IgG sequences (i.e. identical except for constant region) in light blue, IgG/IgA sequences in royal blue, one IgA sequence in red, and reference mAb B cell sequence INF9 shown in gray. White dots indicate putative sequences not found. INF87, found towards the bottom of the lineage and shown in gray, is another memory B cell sequence cloned after HA^+^ FACS but not tested to confirm HA-binding. The sizes of the circles indicate the relative UMI count per sequence. The sequences in this lineage were taken from two replicates of the stimulated PBMC repertoire from the May 2018 timepoint of donor 536. This lineage was made using the Alakazam R scripts in the Immcantation pipeline, and is made using parsimony-based approach, with numbers between sequences indicating the number of mutational steps. **B.** An alignment of variable heavy chain sequences selected from the clonal family lineage shown in A. Somatic hypermutation from germline in the CDR1, CDR2 and CDR3 regions are shown in red and in the framework (FR) regions in orange. The reference mAb INF9 is shown on the bottom line, in bold.

**FIGURE 10.**
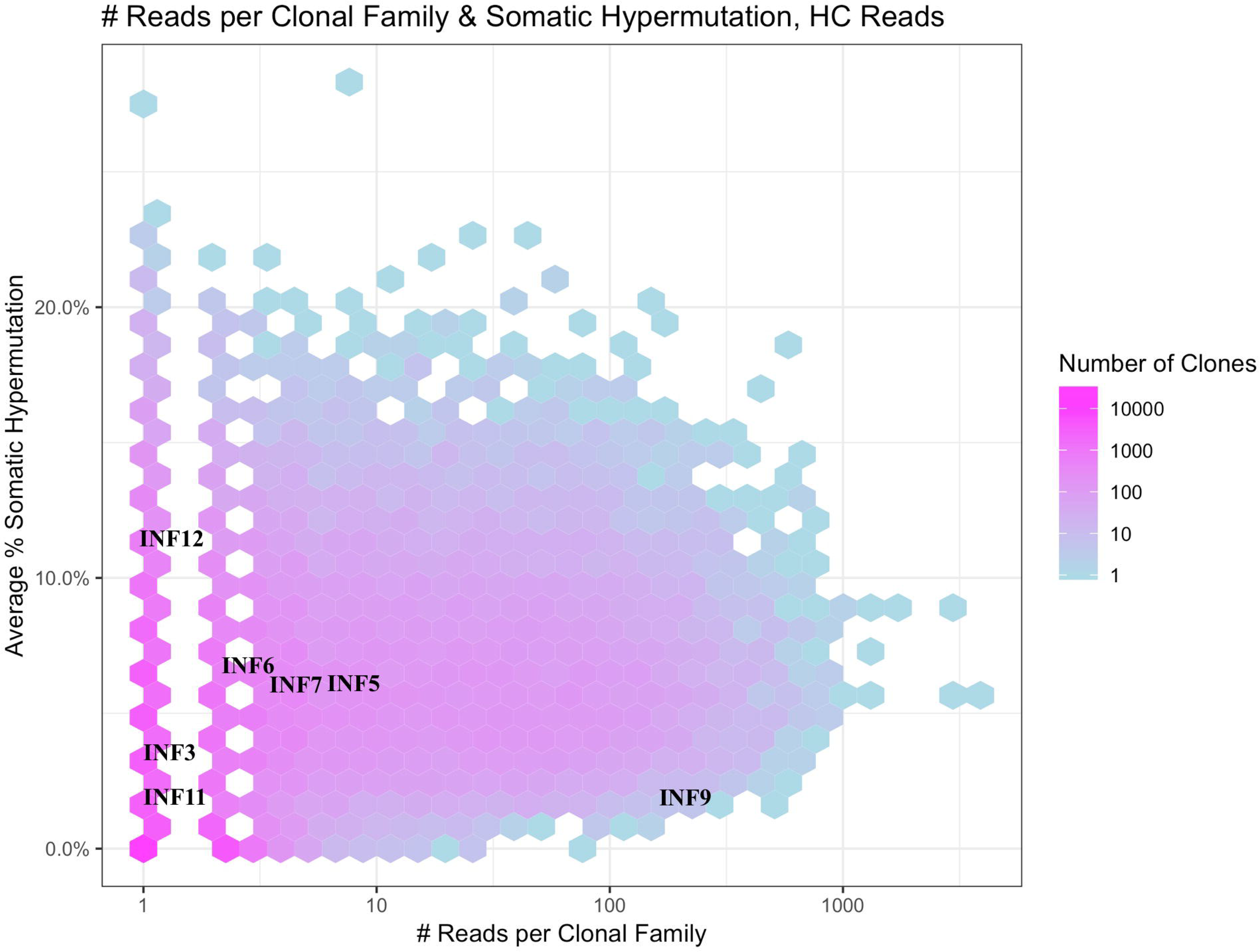
Immunodominance plot of the May stimulated PBMC repertoire of donor 536.

Clonal families of mAb IGVH sequences were found and mapped onto a heat map made from the total May IGVH clonal repertoire. The clonal diversity of the May IGVH repertoire at a given read frequency is indicated using a color scale from pinks for >100 unique clones into blues for <10 unique clones. Since the heat map plots all clones from the repertoire at a given read frequency, individual mAb clonal family diversity cannot be interpreted by color. The y-axis shows the average somatic hypermutation of clonal families and the x-axis the abundance of clonal families by number of sequencing reads. Because whole numbers are plotted using a log scale on the x-axis, this gives rise to an appearance of gaps between the lowest values. The plot was made using the Alakazam R scripts in the Immcantation pipeline.

## DISCUSSION

There is an ongoing need for the development of improved vaccines to replace those that have high production costs, poor stability, efficacy and/or durability as well as a need for research to inform new vaccines for intractable or emerging pathogens. Even for the most successful vaccines, continued vaccination of global populations and surveillance is often necessary to offset the re-emergence of diseases after incomplete geographical or lifespan coverage of immune protection. Furthermore, the natural evolution of pathogens to new or vaccine resistant strains provides an ongoing challenge to protective strategies. An ideal vaccine would provide robust, directed immunogenicity to give rise to dominant clones of high affinity antibodies with specificity, cross-reactivity and pathogen neutralizing or clearing activity. Importantly, clonal families for the corresponding antibody producing B cells would be found to be consistently well represented in the memory compartment, with the ability to be rapidly and functionally activated upon subsequent exposures.

In this paper we provide methods to allow for a deep analysis of the clonal diversity, persistence and hierarchy of antibodies in the memory repertoire from PBMCs. Calculations using the Immcantation pipeline on our stimulated PBMC NGS data provide a means to estimate the number of clonal families attributed to human memory B cells in blood. We observed in the range of six to twenty thousand clonal families for IgG memory among the three donors, but additional data analysis is warranted to capture the variability between replicates and statistical saturation point. Previous work has estimated the plasma cell derived clonal repertoire size in serum to be on the order of 20,000 (Wine Y. *et al* 2015). In future studies, it will be of interest to compare the repertoire of polyclonal *in vitro* stimulation of B cell memory with that of antigen-specific stimulation, and to use that data to identify public (many donor) versus private (individual donor) sequences. Overall, the analysis of deep functional memory repertoires will provide for a better view of the *in vivo* landscape of humoral protection or enhancement of disease, informative to vaccine and therapeutic antibody research.

Clonal expansions of B cells occur in response to specific immune stimulation and evidence for a corresponding convergence of germline gene arrangements has been shown (e.g. Briney B. *et al* 2019, Setliff, I. *et al* 2018, Imkeller, K.,and Wardemann, H. 2018). Clonal families represent *in vivo* antibody molecular evolution to a consensus directed against common epitopes of vaccines or disease. As the field gains more and more knowledge on immune repertoires and collects data on responses to disease and vaccination, we will get closer to being able to deliberately direct our immune system towards epitopes yielding positive health outcomes.

## AUTHOR CONTRIBUTIONS

KM conceived and designed the study. EW and KM carried out the experiments, performed data analysis and wrote the manuscript. EW and AM developed the sequence analysis pipeline. PK provided study oversight and review. All the authors read and approved the final manuscript.

### ACKNOWLDGEMENTS

We thank Payton Anders Weidenbacher and Natalia Friedland of Stanford University, for providing the hemagglutinin and MEDI8852 proteins used for single cell antibody FACS and characterization. We also thankfully acknowledge the expertise in our CZ Biohub genomics group including Norma Neff, Michelle Tan, Rene Sit, and Brian Yu.

## Conflict of Interest Statement

The authors declare that the research was conducted without any influence from commercial or financial relationships.

