## Supplemental Tables 1-3 for "Functional Enrichment and Analysis of Antigen-Specific Memory B Cell Antibody Repertoires in PBMCs"

### SUPPLEMENTARY TABLES

#### Table S1. Amplicon Primers.

| | |
|---|---|
| IgH_constant RT primer pool[1] | |
| RT_IgA_08N | TGACTGGAGTTCAGACGTGTGCTCTTCCGATCT(N:25252525)<br>(N)(N)(N)(N)(N)(N)(N)GGGGAAGAAGCCCTGGAC |
| RT_IgA_12N | TGACTGGAGTTCAGACGTGTGCTCTTCCGATCT(N:25252525)<br>(N)(N)(N)(N)(N)(N)(N)(N)(N)(N)(N)GGGGAAGAAGCCCTGGAC |
| RT_IgG_08N | TGACTGGAGTTCAGACGTGTGCTCTTCCGATCT(N:25252525)<br>(N)(N)(N)(N)(N)(N)(N)GGGAAGTAGTCCTTGACCA |
| RT_IgG_12N | TGACTGGAGTTCAGACGTGTGCTCTTCCGATCT(N:25252525)<br>(N)(N)(N)(N)(N)(N)(N)(N)(N)(N)(N)GGGAAGTAGTCCTTGACCA |
| RT_IgM_long_8N | TGACTGGAGTTCAGACGTGTGCTCTTCCGATCT(N:25252525)<br>(N)(N)(N)(N)(N)(N)(N)GAAGGAAGTCCTGTGCGAG |
| RT_IgM_long_12N | TGACTGGAGTTCAGACGTGTGCTCTTCCGATCTN(N:25252525)<br>(N)(N)(N)(N)(N)(N)(N)(N)(N)(N)(N)(N)GAAGGAAGTCCTGTGCGAG |
| RT_IgE_long_8N | TGACTGGAGTTCAGACGTGTGCTCTTCCGATCT(N:25252525)<br>(N)(N)(N)(N)(N)(N)(N)AAGTAGCCCGTGGCCAGG |
| RT_IgE_long_12N | TGACTGGAGTTCAGACGTGTGCTCTTCCGATCT(N:25252525)<br>(N)(N)(N)(N)(N)(N)(N)(N)(N)(N)(N)AAGTAGCCCGTGGCCAGG |
| RT_IgD_long_8N | TGACTGGAGTTCAGACGTGTGCTCTTCCGATCT(N:25252525)<br>(N)(N)(N)(N)(N)(N)(N)TGGGTGGTACCCAGTTATCAA |
| RT_IgD_long_12N | TGACTGGAGTTCAGACGTGTGCTCTTCCGATCT(N:25252525)<br>(N)(N)(N)(N)(N)(N)(N)(N)(N)(N)(N)TGGGTGGTACCCAGTTATCAA |
| LC_constant RT primer pool[1] | |
| kappa.rev_08N | TGACTGGAGTTCAGACGTGTGCTCTTCCGATCT(N:25252525)<br>(N)(N)(N)(N)(N)(N)(N)AGTTCCAGATTTCAACTGCTCATCAGAT |
| kappa.rev_12N | TGACTGGAGTTCAGACGTGTGCTCTTCCGATCT(N:25252525)<br>(N)(N)(N)(N)(N)(N)(N)(N)(N)(N)(N)AGTTCCAGATTTCAACTGCTCATCAGAT |
| lambda.rev_08N | TGACTGGAGTTCAGACGTGTGCTCTTCCGATCT(N:25252525)<br>(N)(N)(N)(N)(N)(N)(N)GAGGGCGGGAACAGAGTGAC |
| lambda.rev_12N | TGACTGGAGTTCAGACGTGTGCTCTTCCGATCT(N:25252525)<br>(N)(N)(N)(N)(N)(N)(N)(N)(N)(N)(N)GAGGGCGGGAACAGAGTGAC |
| IgH_V forward primer pool[1] | |
| IGH.forP1_08N | ACACTCTTTCCCTACACGACGCTCTTCCGATCT(N:25252525)<br>(N)(N)(N)(N)(N)(N)(N)SCAGCTGGTGCAGTCTGG |
| IGH.forP1_12N | ACACTCTTTCCCTACACGACGCTCTTCCGATCT(N:25252525)<br>(N)(N)(N)(N)(N)(N)(N)(N)(N)(N)(N)SCAGCTGGTGCAGTCTGG |
| IGH.forP135_08N | ACACTCTTTCCCTACACGACGCTCTTCCGATCT(N:25252525)<br>(N)(N)(N)(N)(N)(N)(N)GTGCAGCTGGTGGAGTCTG |
| IGH.forP135_12N | ACACTCTTTCCCTACACGACGCTCTTCCGATCT(N:25252525)<br>(N)(N)(N)(N)(N)(N)(N)(N)(N)(N)(N)GTGCAGCTGGTGGAGTCTG |
| IGH.forP2_08N | ACACTCTTTCCCTACACGACGCTCTTCCGATCT(N:25252525)<br>(N)(N)(N)(N)(N)(N)(N)TCACCTTGAAGGAGTCTGG |
| IGH.forP2_12N | ACACTCTTTCCCTACACGACGCTCTTCCGATCT(N:25252525)<br>(N)(N)(N)(N)(N)(N)(N)(N)(N)(N)(N)TCACCTTGAAGGAGTCTGG |
| IGH.forP4.1_08N | ACACTCTTTCCCTACACGACGCTCTTCCGATCT(N:25252525)<br>(N)(N)(N)(N)(N)(N)(N)TGCAGCTGCAGGAGTCG |
| IGH.forP4.1_12N | ACACTCTTTCCCTACACGACGCTCTTCCGATCT(N:25252525)<br>(N)(N)(N)(N)(N)(N)(N)(N)(N)(N)(N)TGCAGCTGCAGGAGTCG |
| IGH.forP4.2_08N | ACACTCTTTCCCTACACGACGCTCTTCCGATCT(N:25252525)<br>(N)(N)(N)(N)(N)(N)(N)GTGCAGCTACAGCAGTGG |
| IGH.forP4.2_12N | ACACTCTTTCCCTACACGACGCTCTTCCGATCT(N:25252525)<br>(N)(N)(N)(N)(N)(N)(N)(N)(N)(N)(N)GTGCAGCTACAGCAGTGG |
| IGH.forP6_08N | ACACTCTTTCCCTACACGACGCTCTTCCGATCT(N:25252525) |

| | |
|---|---|
| | (N)(N)(N)(N)(N)(N)(N)GTACAGCTGCAGCAGTCA |
| IGH.forP6_12N | ACACTCTTTCCCTACACGACGCTCTTCCGATCT(N:25252525)(N)(N)(N)(N)(N)(N)(N)(N)(N)(N)(N)GTACAGCTGCAGCAGTCA |
| LC_V forward primer pool[1] | |
| Vka_08N | ACACTCTTTCCCTACACGACGCTCTTCCGATCT(N:25252525)(N)(N)(N)(N)(N)(N)(N)GACATCCRGDTGACCCAGTCTCC |
| Vka_12N | ACACTCTTTCCCTACACGACGCTCTTCCGATCT(N:25252525)(N)(N)(N)(N)(N)(N)(N)(N)(N)(N)(N)GACATCCRGDTGACCCAGTCTCC |
| VKb_08N | ACACTCTTTCCCTACACGACGCTCTTCCGATCT(N:25252525)(N)(N)(N)(N)(N)(N)(N)GAAATTGTRWTGACRCAGTCTCC |
| VKb_12N | ACACTCTTTCCCTACACGACGCTCTTCCGATCT(N:25252525)(N)(N)(N)(N)(N)(N)(N)(N)(N)(N)(N)GAAATTGTRWTGACRCAGTCTCC |
| VKc_08N | ACACTCTTTCCCTACACGACGCTCTTCCGATCT(N:25252525)(N)(N)(N)(N)(N)(N)(N)GATATTGTGMTGACBCAGWCTCC |
| VKc_12N | ACACTCTTTCCCTACACGACGCTCTTCCGATCT(N:25252525)(N)(N)(N)(N)(N)(N)(N)(N)(N)(N)(N)GATATTGTGMTGACBCAGWCTCC |
| VKd_08N | ACACTCTTTCCCTACACGACGCTCTTCCGATCT(N:25252525)(N)(N)(N)(N)(N)(N)(N)GAAACGACACTCACGCAGTCTC |
| VKd_12N | ACACTCTTTCCCTACACGACGCTCTTCCGATCT(N:25252525)(N)(N)(N)(N)(N)(N)(N)(N)(N)(N)(N)GAAACGACACTCACGCAGTCTC |
| Vla_08N | ACACTCTTTCCCTACACGACGCTCTTCCGATCT(N:25252525)(N)(N)(N)(N)(N)(N)(N)CAGTCTGTSBTGACGCAGCCGCC |
| Vla_12N | ACACTCTTTCCCTACACGACGCTCTTCCGATCT(N:25252525)(N)(N)(N)(N)(N)(N)(N)(N)(N)(N)(N)CAGTCTGTSBTGACGCAGCCGCC |
| VLb_08N | ACACTCTTTCCCTACACGACGCTCTTCCGATCT(N:25252525)(N)(N)(N)(N)(N)(N)(N)TCCTATGWGCTGACWCAGCCAC |
| VLb_12N | ACACTCTTTCCCTACACGACGCTCTTCCGATCT(N:25252525)(N)(N)(N)(N)(N)(N)(N)(N)(N)(N)(N)TCCTATGWGCTGACWCAGCCAC |
| VLc_08N | ACACTCTTTCCCTACACGACGCTCTTCCGATCT(N:25252525)(N)(N)(N)(N)(N)(N)(N)TCCTATGAGCTGAYRCAGCYACC |
| VLc_12N | ACACTCTTTCCCTACACGACGCTCTTCCGATCT(N:25252525)(N)(N)(N)(N)(N)(N)(N)(N)(N)(N)(N)TCCTATGAGCTGAYRCAGCYACC |
| VLd_08N | ACACTCTTTCCCTACACGACGCTCTTCCGATCT(N:25252525)(N)(N)(N)(N)(N)(N)(N)CAGCCTGTGCTGACTCARYC |
| VLd_12N | ACACTCTTTCCCTACACGACGCTCTTCCGATCT(N:25252525)(N)(N)(N)(N)(N)(N)(N)(N)(N)(N)(N)CAGCCTGTGCTGACTCARYC |
| Vle_08N | ACACTCTTTCCCTACACGACGCTCTTCCGATCT(N:25252525)(N)(N)(N)(N)(N)(N)(N)CAGDCTGTGGTGACYCAGGAGCC |
| Vle_12N | ACACTCTTTCCCTACACGACGCTCTTCCGATCT(N:25252525)(N)(N)(N)(N)(N)(N)(N)(N)(N)(N)(N)CAGDCTGTGGTGACYCAGGAGCC |
| VLf_08N | ACACTCTTTCCCTACACGACGCTCTTCCGATCT(N:25252525)(N)(N)(N)(N)(N)(N)(N)CAGCCWGKGCTGACTCAGCCMCC |
| VLf_12N | ACACTCTTTCCCTACACGACGCTCTTCCGATCT(N:25252525)(N)(N)(N)(N)(N)(N)(N)(N)(N)(N)(N)CAGCCWGKGCTGACTCAGCCMCC |
| VLg_08N | ACACTCTTTCCCTACACGACGCTCTTCCGATCT(N:25252525)(N)(N)(N)(N)(N)(N)(N)TCCTCTGAGCTGASTCAGGASCC |
| VLg_12N | ACACTCTTTCCCTACACGACGCTCTTCCGATCT(N:25252525)(N)(N)(N)(N)(N)(N)(N)(N)(N)(N)(N)TCCTCTGAGCTGASTCAGGASCC |
| VLh_08N | ACACTCTTTCCCTACACGACGCTCTTCCGATCT(N:25252525)(N)(N)(N)(N)(N)(N)(N)CAGTCTGYYCTGAYTCAGCCT |
| VLh_12N | ACACTCTTTCCCTACACGACGCTCTTCCGATCT(N:25252525)(N)(N)(N)(N)(N)(N)(N)(N)(N)(N)(N)CAGTCTGYYCTGAYTCAGCCT |
| Vli_08N | ACACTCTTTCCCTACACGACGCTCTTCCGATCT(N:25252525)(N)(N)(N)(N)(N)(N)(N)AATTTTATGCTGACTCAGCCCC |
| Vli_12N | ACACTCTTTCCCTACACGACGCTCTTCCGATCT(N:25252525)(N)(N)(N)(N)(N)(N)(N)(N)(N)(N)(N)AATTTTATGCTGACTCAGCCCC |
| Illumina sample index primers[2] | |

| | |
|---|---|
| PE1_A6 | AAT GAT ACG GCG ACC ACC GAG ATC TAC ACC GGT TAA AAC ACT CTT TCC CTA CAC GAC GCT CTT CCG ATC T |
| PE2_A6 | CAA GCA GAA GAC GGC ATA CGA GAT AAA TTG GCG TGA CTG GAG TTC AGA CGT GTG CTC TTC CGA TCT |
| PE1_A12 | AAT GAT ACG GCG ACC ACC GAG ATC TAC ACG AAC ATA AAC ACT CTT TCC CTA CAC GAC GCT CTT CCG ATC T |
| PE2_A12 | CAA GCA GAA GAC GGC ATA CGA GAT AAT ACA AGG TGA CTG GAG TTC AGA CGT GTG CTC TTC CGA TCT |
| PE1_A4 | AAT GAT ACG GCG ACC ACC GAG ATC TAC ACA CTG GTA AAC ACT CTT TCC CTA CAC GAC GCT CTT CCG ATC T |
| PE2_A4 | CAA GCA GAA GAC GGC ATA CGA GAT AAT GGT CAG TGA CTG GAG TTC AGA CGT GTG CTC TTC CGA TCT |

[1]The 5' end of the primers correspond to adapter sequences for PE indexing, the middle 8 or 12 Ns for random barcode UMIs (unique molecular identifiers), followed by gene-specific sequences (constant domain reverse or framework 1 region forward). Primer design was based on primers used in Horns F. et al 2016 and Vollmers C. et al 2013.

[2]Refer to Illumina guidelines on PE primer multiplexing (e.g. A1 does not multiplex well with A2 and A3 when the sample number is less than 5).

#### TABLE S2. Antibodies used for Single Memory B Cell FACS.

| Marker | Channel | Supplier | Clone/Product# |
|---|---|---|---|
| CD19-BV421 | FL1 | BioLegend | SJ25C1 |
| SYTOX-Green | FL2 | Thermo Fisher Scientific | S34860 |
| CD3-FITC | FL2 | BioLegend | OKT3 |
| CD14-FITC | FL2 | BioLegend | M5E2S |
| CD56-FITC | FL2 | BD Biosciences | B159 |
| Anti-human IgD-AlexaFluor 488 | FL2 | BioLegend | 1A6-2 |
| Anti-human IgM-FITC | FL2 | BioLegend | MHM-88 |
| Anti-mouse IgA-FITC | FL2 | BD Biosciences | C10-3 |
| CD27-PE | FL3 | BioLegend | M-T271 |
| Streptavidin-AlexaFluor 647 | FL4 | BioLegend | 405237 |
| CD20-PECy7 | FL6 | BioLegend | 2H7 |

**TABLE S3. Primer Sequences used for Human Antibody Cloning.**

| HEAVY IgG CHAIN | | 100uM stock |
|---|---|---|
| VHa | CAGGTGCAGCTGCAGGAGTCSG | 10ul |
| VHb | CAGGTACAGCTGCAGCAGTCA | 10ul |
| VHc | CAGGTGCAGCTACAGCAGTGGG | 10ul |
| VHd | GAGGTGCAGCTGKTGGAGWCY | 10ul |
| VHe | CAGGTCCAGCTKGTRCAGTCTGG | 10ul |
| VHf | CAGRTCACCTTGAAGGAGTCTG | 10ul |
| VHg | CAGGTGCAGCTGGTGSARTCTGG | 10ul |
| Water | | 15ul |
| IGG.hingeU.reverse | CTGGGCAYSRTGGGCAY | 15ul |
| Total | | 100ul |
| LIGHT CHAIN (KAPPA) | | |
| Vka | GACATCCRGDTGACCCAGTCTCC | 10ul |
| VKb | GAAATTGTRWTGACRCAGTCTCC | 10ul |
| VKc | GATATTGTGMTGACBCAGWCTCC | 10ul |
| VKd | GAAACGACACTCACGCAGTCTC | 10ul |
| Water | | 45ul |
| CK.reverse | ACTAACACTCTCCCCTGTTGAAGCTCTTTGTGACGGG CGATCTCA | 15ul |
| Total | | 100ul |
| LIGHT CHAIN (LAMBDA) | | |
| Vla | CAGTCTGTSBTGACGCAGCCGCC | 10ul |
| VLb | TCCTATGWGCTGACWCAGCCAC | 10ul |
| VLc | TCCTATGAGCTGAYRCAGCYACC | 10ul |
| VLd | CAGCCTGTGCTGACTCARYC | 10ul |
| Vle | CAGDCTGTGGTGACYCAGGAGCC | 10ul |
| VLf | CAGCCWGKGCTGACTCAGCCMCC | 10ul |
| VLg | TCCTCTGAGCTGASTCAGGASCC | 10ul |
| VLh | CAGTCTGYYCTGAYTCAGCCT | 10ul |
| Vli | AATTTTATGCTGACTCAGCCCC | 10ul |
| CL.reverse | ATCTGCCTTCCAGGCCACTGTCAC | 15ul |
| Total | | 105ul |
| SEQUENCING PRIMERS | | |
| IGG.CH1.rev | GGGAAGTAGTCCTTGACCA | |
| kappa.rev | AGTTCCAGATTTCAACTGCTCATCAGAT | |
| lambda.rev | AGAGGGCGGGAACAGAGTGAC | |
